# Synaptonemal complex-deficient *Drosophila melanogaster* females exhibit rare DSB repair events, recurrent copy number variation, and an increased rate of *de novo* transposable element movement

**DOI:** 10.1101/814186

**Authors:** Danny E. Miller

## Abstract

Genetic stability depends on the maintenance of a variety of chromosome structures and the precise repair of DNA breaks. During meiosis, programmed double-strand breaks (DSBs) made in prophase I are normally repaired as gene conversions or crossovers. Additionally, DSBs are made by the movement of transposable elements (TEs), which must also be resolved. Incorrect repair of these DNA lesions can lead to mutations, copy number variations, translocations, and/or aneuploid gametes. In *Drosophila melanogaster*, as in most organisms, meiotic DSB repair occurs in the presence of a rapidly evolving multiprotein structure called the synaptonemal complex (SC). Here, whole-genome sequencing is used to investigate the fate of meiotic DSBs in *D. melanogaster* mutant females lacking functional SC, to assay for de novo CNV formation, and to examine the role of the SC in transposable element movement in flies. The data indicate that, in the absence of SC, copy number variation still occurs but meiotic DSB repair by gene conversion may occur only rarely. Remarkably, an 856-kilobase de novo CNV was observed in two unrelated individuals of different genetic backgrounds and was identical to a CNV recovered in a previous wild-type study, suggesting that recurrent formation of large CNVs occurs in Drosophila. In addition, the rate of novel TE insertion was markedly higher than wild type in one of two SC mutants tested, suggesting that SC proteins may contribute to the regulation of TE movement and insertion in the genome. Overall, this study provides novel insight into the role that the SC plays in genome stability and provides clues as to why SC proteins are among the most rapidly evolving in any organism.

## INTRODUCTION

Programmed double-stranded DNA breaks (DSBs) made during prophase of meiosis I are a critical step in the formation of healthy gametes, yet they are potentially catastrophic events for cells. The meiotic break repair machinery must therefore accurately resolve DSBs as either crossovers (COs) or noncrossover gene conversions (NCOGCs). More DSBs are made than will be repaired as COs, and thus the majority of DSBs are repaired as NCOGCs which are nonreciprocal exchange events that result in the 3:1 segregation of alleles. Crossover-associated gene conversions—those that occur in conjunction with a crossover—are frequently seen in some organisms (Jeffreys and May 2004; Santoyo *et al.* 2005; Mancera *et al.* 2008; Wijnker *et al.* 2013), but are less frequently observed in Drosophila (Curtis *et al.* 1989; Hilliker *et al.* 1994; Miller *et al.* 2016).

Crossing over is essential to ensure the proper segregation of homologous chromosomes during the subsequent meiotic divisions. Crossing over occurs within the context of a large multiprotein structure called the synaptonemal complex (SC), which forms between homologous chromosomes. In most organisms, DSBs must be made before SC formation can occur, and functional SC is required for proper DSB repair (de Massy 2012; Zickler and Kleckner 2015). However, in *Drosophila melanogaster* the SC is necessary for both robust DSB formation and DSB repair (Lake and Hawley 2012); in the absence of functional SC, DSBs are made at about 20–40% of the wild-type level (Mehrotra and McKim 2005; Collins *et al.* 2014).

The Drosophila SC protein C(3)G is functionally homologous to the transverse filament proteins SYCP-1 in mammals and ZIP1 in budding yeast (Page and Hawley 2001). While females heterozygous for a loss-of-function *c(3)G* allele appear to build normal SC, homozygous females do not build SC and are thus unable to resolve into crossovers those DSBs that do occur (Page and Hawley 2001). A previous study examining NCOGC events at a single locus in Drosophila recovered no events from *c(3)G* homozygous females but did not report the number of progeny scored (Carlson 1972), thus whether DSBs can be repaired as NCOGCs in females lacking functional SC is unknown. Like C(3)G, the Drosophila SC protein Corolla is also required for SC formation. *corolla* mutants exhibit phenotypes typical of Drosophila SC mutants, including a reduced number of DSBs (∼40% as assayed by γH2AV foci) and increased levels of chromosome segregation defects (Collins *et al.* 2014). Similar to *c(3)G* homozygous females, how DSBs are repaired in *corolla* homozygotes remains unknown.

While it is evident the SC plays a vital role in resolving DSBs into COs, its role in other meiotic processes is less obvious. For example, there is some evidence for a link between SC formation and transposable elements (TEs), but the data are not definitive (Pearlman *et al.* 1992; Hernández-Hernández *et al.* 2008; Marcon *et al.* 2008; van der Heijden and Bortvin 2009). Transposable elements are mobile genetic elements active during different stages of gametogenesis. They can be divided into two classes: Class 1, or retrotransposons, replicate using a copy-and-paste method to insert copies of themselves into new locations in the genome, while Class 2, or DNA transposons, use a cut-and-paste method to move from one position in the genome to another. SC genes are among the most rapidly evolving genes in all organisms, but the reason for this remains unknown. It has been hypothesized that the rapid evolution of SC genes may occur to counter the effects of transposable element (TE) movement during meiosis (Fraune *et al.* 2012; Hemmer and Blumenstiel 2016). In Drosophila female meiosis, the rate at which TE movement occurs and if the SC has any role in facilitating or limiting TE movement remains unclear.

In the current study, whole-genome sequencing (WGS) was used to investigate individual meiotic events in male offspring from females heterozygous or homozygous for a loss-of-function allele of *c(3)G*. While the number and distribution of CO and NCOGC events in individuals from females heterozygous for *c(3)G* was similar to wild-type, in progeny arising from *c(3)G* homozygous mothers (which do not build SC), no crossovers and only one likely NCOGC event were recovered. The recovery of a single presumed NCOGC event suggests that while repair of DSBs via NCOGC may be possible in females lacking functional SC, it is extremely rare. Consistent with the high levels of chromosome missegregation observed in SC mutants, *X0* males lacking a *Y* chromosome, males with *4*^*th*^ chromosome gain or loss, and intersex males were also recovered.

These data also provide information on what role, if any, the SC components C(3)G and Corolla play in facilitating or inhibiting TE movement during meiosis. In the current study of SC-defective mutants, novel TE insertions were curiously significantly elevated in *c(3)G* homozygotes but similar to wild type in *corolla* homozygotes. Previous work observed an unexpectedly high amount of transposable element (TE)-mediated copy number variation (CNV) between sister chromatids in wild-type *Drosophila* offspring (Miller *et al.* 2016). Shared and novel large-scale TE-mediated CNVs were also identified in progeny from all genotypes. Remarkably, one of these CNVs was observed in three unrelated individuals—two from this study and one from a separate study of individual meiotic events in wild type (Miller *et al.* 2016)—suggesting that, similar to humans (Itsara *et al.* 2009), recurrent CNVs may be a common occurrence in Drosophila. Overall, this work helps further our understanding of how meiotic cells cope with DNA breaks and maintain genetic stability.

## METHODS

### Fly Stocks and husbandry

The loss-of-function allele *c(3)G*^*68*^ (Page and Hawley 2001) was crossed into stocks isogenic for either *w*^*1118*^ or Canton-S strain polymorphisms (Miller *et al.* 2012). Females homozygous for *Canton-S X* and *2*^*nd*^ chromosomes and heterozygous for the *c(3)G*^*68*^ loss-of-function allele were crossed to *w*^*1118*^ males to generate females heterozygous for Canton-S and *w*^*1118*^ strain polymorphisms. These heterozygous females were then crossed again to isogenic *w*^*1118*^ males and individual male progeny were isolated for sequencing (Figure S1). Females heterozygous for *w*^*1118*^ and Canton-S *X* and *2*^*nd*^ chromosomes and homozygous for *c(3)G*^*68*^ were crossed to isogenic *w*^*1118*^ males and individual male offspring were isolated for sequencing (Figure S1). Progeny from *corolla*^*129*^ homozygous females were generated by crossing virgin *corolla*^*129*^ females to sibling males and collecting both male and female progeny (Figure S1). All crosses were done using a single male and female, and females were allowed to lay eggs for 7 days before being removed from a vial. Male offspring used for sequencing were collected between days 12 and 15. All flies were kept on standard cornmeal-molasses and maintained at 25°C.

### DNA preparation and sequencing

For all flies, DNA was prepared from single adult males or females using the Qiagen DNeasy Blood & Tissue Kit. All flies were starved for 4 hr before freezing at −80°C for at least 1 hr. One µg of DNA from each was fragmented to 250-bp fragments by adjusting the treatment time to 85 sec using a Covaris S220 sonicator (Covaris Inc.). Libraries were prepared using a Nextera DNA Sample Prep Kit and Bioo Scientific NEXTflex DNA Barcodes. The resulting libraries were purified using Agencourt AMPure XP system (Beckman Coulter) then quantified using a Bioanalyzer (Agilent Technologies) and a Qubit Fluorometer (Life Technologies). Samples from *c(3)G*^*68*^ homozygotes females were run on a HiSeq 2500 in rapid mode as either 100-bp paired-end or 125-bp paired-end samples using HiSeq Control Software 1.8.2 and Real-Time Analysis (RTA) version 1.17.21.3. Samples from the *c(3)G*^*68*^ heterozygous and *corolla*^*129*^ homozygous experiments were run as 150-bp paired-end on a HiSeq 2500 in rapid mode using HiSeq Control Software 2.2.58 and RTA version 1.18.64. Secondary Analysis version CASAVA-1.8.2 was run to demultiplex reads and generate FASTQ files. Per-sample sequencing and alignment statistics can be found in Table S1.

### DNA alignment, SNP calling, and identification of CO and NCOGC events

Alignment to the Drosophila reference genome (dm6) was preformed using bwa version 0.7.7-r441 using default paramaters (Li and Durbin 2009). Single nucleotide and insertion or deletion polymorphisms were identified using SAMtools version 1.9 (Li *et al.* 2009). Candidate CO and NCOGC events were identified as described in Miller *et al*. (Miller *et al.* 2016).

### Depth-of-coverage calculations

Depth of coverage for each chromosome arm was calculated by summing the total read depth for each base position then dividing by the length of the entire chromosome arm. Because of the repetitive nature of the *Y* chromosome, analysis was limited to *chrY*:332,000–510,000 (Table S1).

### Validation of NCOGCs by PCR

Nine candidate NCOGC events were identified in 93 males from *c(3)G*^*68*^ females and examined by PCR and Sanger sequencing; Phusion polymerase (NEB) was used according to the manufacturer’s instructions. Only one of the nine putative conversion events validated as real in male c3g6.4. All primers used can be found in Table S2.

### Calculation of expected NCOGC events

The number of NCOGCs expected to be recovered from 93 individuals from *c(3)G*^*68*^ females if all DSBs on the *X* and *2*^*nd*^ chromosomes were repaired as NCOGCs was estimated by performing 100,000 trials of randomly distributing an estimated number of DSBs among the *X* and *2*^*nd*^ chromosomes using the SNP density of a *w*^*1118*^/*Canton-S* heterozygote. Given that DSBs in *c(3)G*^*68*^ females are made at 20% of the wild-type rate of 18–20 DSBs per meiosis (Mehrotra and McKim 2005), a per-arm number of DSBs was estimated as 0–2 per meiosis. Each break was randomly assigned to a chromosome arm, then to a random chromatid. A random chromatid was then selected to be recovered. NCOGC tract length was assumed to be a minimum of 250 bp and a maximum of 1000 bp (Miller *et al.* 2016). An NCOGC was predicted to be recoverable if the tract involved at least one high-quality SNP that differentiated the *w*^*1118*^ and *Canton-S* genotypes. The estimate of the number NCOGCs which should be recovered from individual offspring of *c(3)G*^*68*^/+ females was calculated by multiplying the wild-type per-arm NCOGC rate of 0.3 (Miller *et al.* 2016) by 120, the number of arms studied.

### Identification of novel deletion polymorphisms

Novel deletions were identified using two approaches. Deletions smaller than 30 bp were identified using SAMtools (Li *et al.* 2009). For each class of progeny (wild type, *c(3)G*^*68*^, *c(3)G*^*68*^/+, and *corolla*^*129*^) a custom script identified any deletion, regardless of quality score, from all vcf files that did not overlap repetitive regions as defined by Repeatmasker (AFA *et al.*). Novel deletions were those with quality scores over 200 (as determined by SAMtools) that did not fall within 100 bp of another deletion on a different offspring. Candidate novel deletions were validated visually using IGV (Thorvaldsdóttir *et al.* 2013). Data for both wild-type and *c(3)G*^*68*^ were also analyzed using GATK HaplotypeCaller (McKenna *et al.* 2010), but no deletions not identified by SAMtools were identified, thus the remainder of the analysis was completed with SAMtools. Larger deletions were identified using Pindel (Ye *et al.* 2009). For each class of progeny, Pindel was run using default settings with an average insert size of 200 bp. Output files for each class of progeny were analyzed as a group and candidate novel deletions were visually validated using IGV.

### Construction of synthetic genomes and sequencing reads

In order to determine what percentage of small or large *de novo* deletion polymorphisms would be identified by SAMtools and Pindel synthetic genomes were computationally modified with deletions of varying sizes then analyzed using the approach described above. Two classes of genomes were generated, 100 with 1–10 bp deletions, and 100 with 1–1000 bp deletions. For each individual, two genomes were generated: one with an *X* and without a *Y* chromosome, and one with a *Y* and without an *X*. For each of these genomes, a single nucleotide was randomly changed approximately once every 500 nucleotides to a randomly selected A, G, C, or T. Next, for each genome with an *X* and without a *Y* chromosome 2–6 DSBs (approximately 20% of the 18–20 DSBs expected in wild-type (Mehrotra and McKim 2005)) were randomly placed on one of four haplotypes in a euchromatic location in the genome. Each of these DSBs was randomly determined to have a deletion between either 1–10 bp or 1–1,000 bp beginning at the site of the DSB. One haplotype of these four was then randomly chosen as the genome for the individual. For each individual, ART was used to generate synthetic reads for both genomes with a read depth of approximately 10x (Huang *et al.* 2012). FASTQ files were then combined into a single forward and a single reverse file, and thus represented data from an XY individual, that were then aligned to the *D. melanogaster* reference genome as above. SNPs, insertion/deletion polymorphism, and larger deletions were identified as described above with SAMtools (Li *et al.* 2009)and Pindel (Ye *et al.* 2009). Deletions generated per individual genome can be found in Table S3.

### Identification of transposable element insertions

To identify TE insertions, split and discordant read pairs were isolated from alignment files using SAMBLASTER (Faust and Hall 2014). BLAST (Altschul *et al.* 1997) was then used to annotate individual split or discordant reads using the *D. melanogaster* canonical TE set (Kaminker *et al.* 2002). Split and discordant clusters that contained more than five reads aligning to a specific TE family were considered candidate TE insertion sites. Novel insertions were detected by a custom script that compared insertions in one population or stock to related stocks or populations and were visually validated using IGV (Thorvaldsdóttir *et al.* 2013).

### Identification of CNV events

CNV events were identified as described in Miller *et al*. (Miller *et al.* 2016). Briefly, average depth of coverage for each individual chromosome arm was determined, then the log_2_ depth of coverage for 5-kb nonoverlapping windows was calculated and plotted to reveal large regions of deletions or duplications.

### Data availability

Illumina data generated for this project are available at the National Center for Biotechnology Information (https://www.ncbi.nlm.nih.gov/). Data for males from *c(3)G68* females can be found under project PRJNA565835, data for males from *c(3)G68* heterozygous females is under project PRJNA565834, and data for males and females from corolla129 females is under project PRJNA565794. Wild-type data used in this study were obtained from project PRJNA307070 (Miller et al. 2016). All code used in this project is available at GitHub (https://github.com/danrdanny/c3g-corolla-project/). Supplemental material is available at Figshare.

## RESULTS AND DISCUSSION

### Analysis of individual meiotic events from *c(3)G*^*68*^ heterozygous and homozygous females

While in many organisms DSBs are made in the absence of SC (de Massy 2012; Zickler and Kleckner 2015), Drosophila is unique in that SC is required for robust DSB formation (Lake and Hawley 2012). *D. melanogaster* females homozygous for loss-of-function alleles of SC genes make DSBs at a rate approximately 20%–40% that of wild type (Mehrotra and McKim 2005; Collins *et al.* 2014), and it remains unclear how these DSBs are repaired. Studies using visual markers in Drosophila have shown that repair of DSBs by crossing over is substantially reduced or completely abolished in females unable to construct full-length SC, and it is not known if these DSBs can be repaired by other pathways, such as NCOGC or NHEJ (Gowen 1933; Hall 1972; Page and Hawley 2001; Manheim and McKim 2003; Jeffress *et al.* 2007; Page *et al.* 2007; Collins *et al.* 2014).

To better understand this process, whole-genome sequencing was performed on individual male progeny from mothers heterozygous for wild-type *Canton-S* and *w*^*1118*^ *X* and *2*^*nd*^ chromosomes and either homozygous or heterozygous for the loss-of-function allele *c(3)G*^*68*^ on chromosome *3*. Male progeny from *c(3)G*^*68*^ homozygous mothers (96 males from 10 females) represent the experimental group lacking SC and will hereafter be referred to as *c(3)G* offspring, and male progeny from *c(3)G*^*68*^ heterozygous mothers (40 males from two females) represent the control group with functional SC and will be referred to as *c(3)G/+* offspring (Figure S1).

While Drosophila males normally have an *X* and a *Y* chromosome, sex is determined by the ratio of *X* chromosomes to autosomes rather than the presence of a *Y*, thus *X0* flies are male. This is seen when X chromosome missegregation (nondisjunction) leads to no maternal sex chromosome contribution, with a paternally inherited *X*. Triploid flies carrying three copies of each autosome and two *X* chromosomes are also phenotypically male and are known as intersex males (Bridges 1921). To assay for *X* and *4*^*th*^ chromosome nondisjunction and the presence of triploid flies, depth of coverage was calculated for each chromosome arm as a percentage of one of the autosomes (Table S1). Males carrying the expected number of *X* chromosomes should have *X* and *Y* chromosome depth of coverage half that of an autosome and *4*^*th*^ chromosome depth of coverage equal to an autosome. As expected, all 40 male offspring from the *c(3)G/+* control group were diploid with an *X* and a *Y* chromosome as well as two copies of the *4*^*th*^ chromosome. Meanwhile, among the offspring from the SC-deficient *c(3)G* experimental group 25 were found to be X0 males and thus carried a paternally-inherited *X* chromosome and six were found to have three *4*^*th*^ chromosomes (Table S1, Figure S2). This high level of non-disjunction in *c(3)G* homozygotes was expected and is similar to previous genetic analyses (Hall 1972).

Three of the 96 *c(3)G* offspring had *X* chromosome depth of coverage approximately 67% that of chromosomes *2* and *3*, with two of these three also carrying a *Y* chromosome (Figure S2, Table S1). Allele frequency for each SNP on each chromosome arm was calculated and revealed all three males were triploid, with one XX:222:333 male and two XXY:222:333 males (Figure S3). The XX:222:333 intersex male was also mosaic for loss of a *4*^*th*^ chromosome, with 75% depth of coverage of the *4*^*th*^ compared to chromosome *2L*, suggesting post-meiotic loss of the *4*^*th*^ in XX:222:333:444 cells (Figure S2). The recovery of intersex individuals was not surprising as previous studies have noted an increase in the number of triploid individuals recovered from *c(3)G* mutants (Gowen 1933; Lindsley and Zimm 1992). These three individuals were excluded from subsequent analysis.

CO and NCOGC events were then identified on the *X* and *2*^*nd*^ chromosomes in both *c(3)G* and *c(3)G*/+ male offspring through changes in polymorphisms in each fly. (CO and NCOGC events were not analyzed on the *3*^*rd*^ because *c(3)G* lies on this chromosome nor on the 25 paternally inherited *X* chromosomes carried by *X0 c(3)G* offspring.) A total of 41 single COs and 7 double COs were identified in *c(3)G*/+ offspring (Figure 1A, Table S4), with a frequency of exchange similar to previous observations in wild type for all three arms (Figure 1B). A total of 32 NCOGCs were also identified (Table S5), close to the 36 expected to be recovered based on wild-type rates (Miller *et al.* 2016). Previous work has shown rates of crossing over similar to wild type for the *c(3)G*^*68*^ allele when heterozygous (Hall 1977), but higher rates of crossing over for *c(3)G*^*17*^ as a heterozygote (Hinton 1966). While the *c(3)G*^*68*^ allele is a known point mutation, the *c(3)G*^*17*^ allele (also historically known as *c(3)G*^*1*^) is a transposable element insertion that disrupts the function of the gene (Page and Hawley 2001), and the reason for the difference in exchange between these two alleles is not clear.

**Figure 1.**
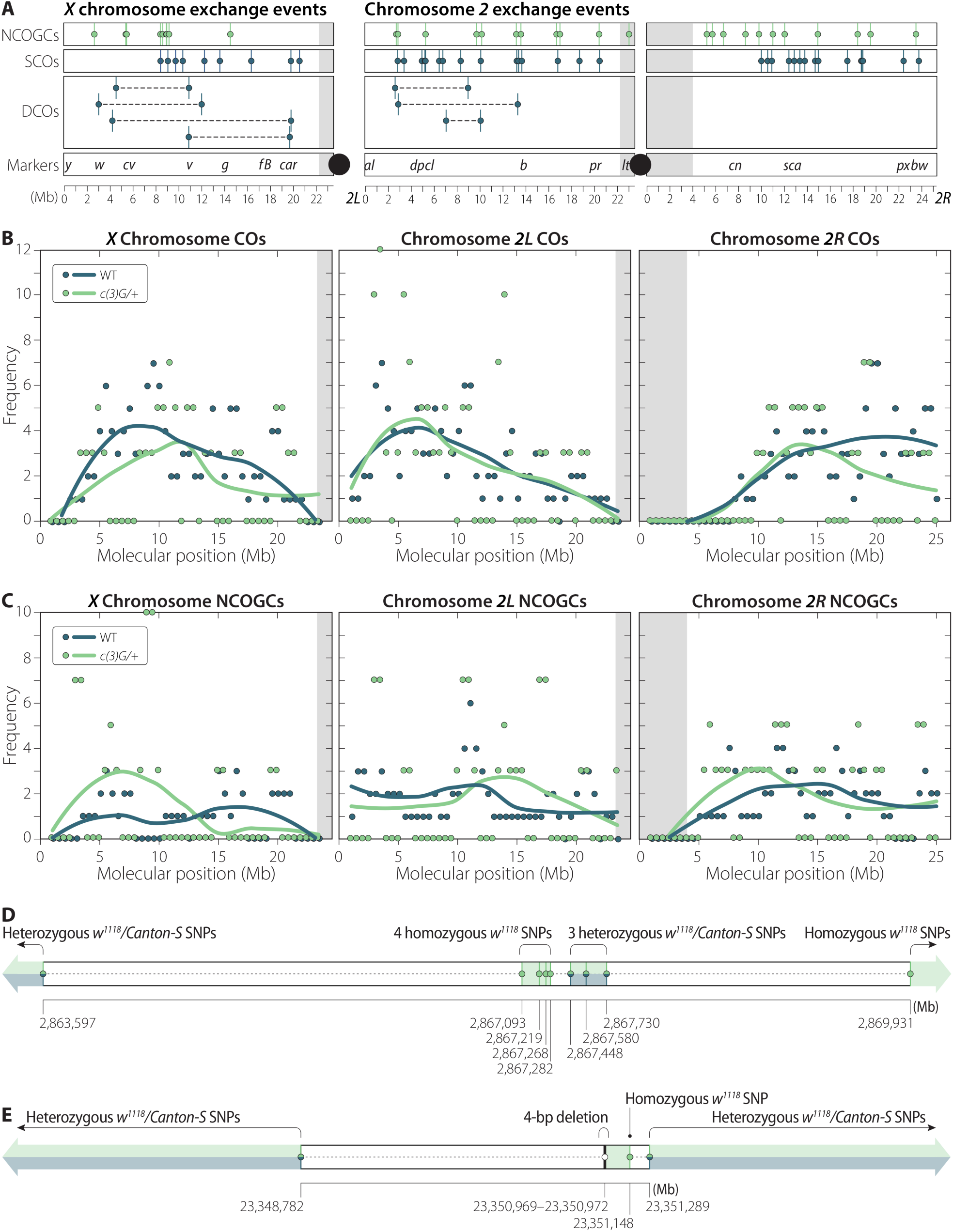
CO and NCOGC events recovered from *c(3)G*^*68*^ heterozygous females, details in Table S4 and Table S5. **A.** individual NCOGC, SCO, and DCO events recovered per chromosome arm. No DCOs were recovered on *2R*. **B.** Coefficient of exchange for all 55 crossover events recovered in this study compared to wild type data from Miller *et al*. (Miller *et al.* 2016). **C.** Coefficient of exchange for all 32 NCOGC events recovered in this study compared to wild type data as in B. **D.** Detail of the single CO-associated GC was recovered in this study. The crossover could have occurred at one of two positions, either between SNPs at positions 2,863,597 and 2,867,093 with the CO-associated GC being the heterozygous tract between positions 2,867,448 and 2,867,730. Alternatively, the CO may have occurred between 2,867,730 and 2,869,931 with the CO-associated GC defined by the 4 SNPs between 2,867,093 and 2,867,282. No CO-associated GC events were recovered in a previous analysis of 196 individual meiotic events from wild-type females (Miller *et al.* 2016). **E.** Structure of the single NCOGC event recovered from a homozygous *c(3)G*^*68*^ female in this study. This NCOGC, validated by PCR and Sanger Sequencing was defined by a 4 bp deletion on one side and a SNP on the other, both from the *w*^*1118*^ line. The NCOGC has a maximum possible tract length of 2,507 bp and a minimum tract length of 180 bp.

Notably, a single crossover-associated gene conversion was identified abutting the distal CO of a double CO in individual c3g-het-3.09 (Figure 1D, Table S4, Table S5). While crossover-associated gene conversions are frequently observed in other organisms (Jeffreys and May 2004; Santoyo *et al.* 2005; Mancera *et al.* 2008; Wijnker *et al.* 2013), a previous study of 196 wild-type meiotic events in *D. melanogaster* found none among 541 CO events, suggesting that these are relatively rare in flies or may be masked due to poor SNP density (Miller *et al.* 2016).

Among 93 *c(3)G* homozygous offspring, no CO events were recovered on the *X* or *2*^*nd*^ chromosomes, but a single NCOGC event in male c3g-hom-6.4 was identified and validated by PCR and Sanger sequencing. This event occurred on a chromosome with the *Canton-S* haplotype, which could have occurred only in the heterozygous *w*^*1118*^/*Canton-S* mother and so was clearly not contributed by the isogenic *w*^*1118*^ father. This NCOGC was minimally defined by a 4-bp deletion on the 5’ side (*2R*:23,350,969–23,350,972, release 6 coordinates) and a single polymorphism on the 3’ side (*2R*:23,351,148) (Figure 1E, Table S5). Because it was defined by these two closely located polymorphisms that created two changes identical to the other haplotype used in this study, it is unlikely the event was the result of *de novo* somatic mutation. The average depth of coverage within the 1-kb interval surrounding the two polymorphisms was 54x, similar to the average depth of coverage for chromosome *2R* for this individual, making it unlikely that this NCOGC was due to a deletion or duplication of this interval. Additionally, the minimum and maximum possible widths of this NCOGC are 180 bp and 2,507 bp, respectively, well within ranges observed in wild type (Bridges 1921). Homologous chromosomes pair prior to meiotic onset, therefore this NCOGC could be the result of DSB repair in a pre-meiotic cell (Bosco 2012; Joyce *et al.* 2012). Unfortunately, there are no reliable estimates of the rate at which this occurs, making the likelihood difficult to assess.

Females homozygous for *c(3)G* loss-of-function alleles make DSBs at ∼20% the level of wild type (Mehrotra and McKim 2005). To estimate the number of NCOGCs that should have been recovered in the *c(3)G* dataset if DSB repair as NCOGCs occurred frequently, a simulation was performed. This model randomly distributed DSBs among 68 *X* and 93 *2*^*nd*^ chromosome arms as if they occurred at a rate 20% that of wild type. This model estimated that 37–62 NCOGCs should have been recovered if all DSBs that occurred were repaired as NCOGCs (since crossovers do not occur in *c(3)G*^*68*^ homozygotes). The recovery of a single candidate NCOGC event is significantly less (*P* < 0.001, Fisher’s exact) than the 37–62 expected NCOGCs, thus repair of DSBs by NCOGC is rare in females unable to construct full-length SC. Therefore, the rate of NCOGC in an SC-deficient female can be estimated as approximately 1×10^−10^ per bp per meiosis, markedly lower than the wild-type rate of 1.9×10^−8^ NCOGCs per bp per meiosis (Hilliker *et al.* 1994; Miller *et al.* 2016). This raises the obvious question, which will be considered next: if DSBs are rarely, if ever, repaired as COs or NCOGCs, what *is* the fate of DSBs that occur in SC-deficient flies?

### DSB repair in SC-deficient females does not result in novel deletion polymorphisms

In addition to meiotic CO or NCOGC, other potential mechanisms for repair of meiotic DSBs exist. Although nonhomologous end-joining (NHEJ) has been described as an error-prone process resulting in deletions (Bétermier *et al.* 2014), data suggest the canonical NHEJ pathway is a higher-fidelity system than previously believed (Kabotyanski *et al.* 1998; Feldmann *et al.* 2000). Alternatively, single-strand annealing (SSA), alternative end-joining (Alt-EJ), and microhomology-mediated end-joining (MMEJ) are pathways that may result in small deletions that could be detected as novel deletion polymorphisms in whole-genome sequencing data (Wang *et al.* 2003, 2005; Guirouilh-Barbat *et al.* 2007; Rass *et al.* 2009). Finally, repair of DSBs using the sister chromatid as a template may occur and leave little or no evidence that could be detected by WGS. Gene conversion with the sister chromatid has been shown to be a significant repair pathway in both *S. cerevisiae* (Goldfarb and Lichten 2010) and mammalian cells (Johnson and Jasin 2000), thus it is reasonable to assume it may be active during Drosophila female meiosis as well. Indeed, ring chromosome assays have shown a decrease in the recovery of ring chromosomes in the absence of *c(3)G*, suggesting breaks in *c(3)G* homozygous females may be repaired by intersister recombination (Sandler 1965). Furthermore, the recovery of *Bar* revertants from *FM7/+*; *c(3)G*^*68*^ females through unequal exchange between sister chromatids supports the hypothesis that sister chromatid exchange occurs in flies as well, although the rate is unknown (Curtis *et al.* 1989; Hilliker *et al.* 1994; Miller *et al.* 2016b).

To determine if DSB repair in SC-deficient females occurs by an error-prone process such as NHEJ, novel deletion polymorphisms were identified in the three previously described classes of progeny (wild-type, *c(3)G*^*68*^ heterozygotes, and *c(3)G*^*68*^ homozygotes) plus an additional class unable to repair DSBs by crossing over. Females carrying loss-of-function mutations of the SC gene *corolla* are unable to construct full-length SC and thus have a high rate of nondisjunction yet still make DSBs at a rate approximately 40% of wild-type (Collins *et al.* 2014), similar to the phenotype observed in *c(3)G* loss-of-function mutations. 50 individual males and females from three females homozygous for a nonsense mutation in the SC protein *corolla* (*corolla*^*129*^) were sequenced. Of these 50 individuals, 11 were the result of *X* chromosome nondisjunction, with 3 *X0* males and 8 *XXY* females; 9 were triplo-4; 2 were nondisjunctional for both the *X* and *4*^*th*^ chromosomes; and no *X* or *4*^*th*^ chromosome mosaics were observed (Table S1). The genetic background of the *2*^*nd*^ and *3*^*rd*^ chromosomes of females homozygous for *corolla*^*129*^ was not controlled, thus candidate CO and NCOGC events could not be identified, but previous studies have shown a nearly complete absence of exchange in *corolla* homozygous females (Collins *et al.* 2014).

*De novo* deletions were searched for using two different approaches (see methods). First, vcf files generated by SAMtools (Li *et al.* 2009) were analyzed for deletion polymorphisms (these are generally less than 20 bp), and second, larger deletions were identified using Pindel (Ye *et al.* 2009). Separately, the output of GATK HaploType caller (McKenna *et al.* 2010)was compared to SAMtools and was found to produce similar results, thus only data from SAMtools was analyzed. Both approaches identified a similar number of *de novo* deletions per fly in all four classes of progeny (wild-type, *c(3)G*^*68*^ heterozygotes, *c(3)G*^*68*^ homozygotes, and *corolla*^*129*^ homozygotes). Specifically, using SAMtools, 11 deletions ranging from 1–11 bp were identified in previously published data from 196 wild-type males, a single 21-bp deletion in 40 *c(3)G*/+ offspring, 8 deletions 1–14 bp large from 93 *c(3)G* offspring, and a single 3 bp deletion in 50 *corolla*^*129*^ individuals were identified (Table S6). Pindel, which searches for larger deletions than would be identified by SAMtools, identified only one novel deletion among all genotypes, a complex 17-bp deletion in a *c(3)G* homozygous male. The recovery of deletions at a rate similar to wild-type suggests that DSBs are repaired by a non-error-prone process with the homolog or with the sister chromatid. However, a caveat of this analysis is that secondary alignment and/or analysis errors make these events difficult to detect.

To test whether the analysis approach was robust enough to detect both large and small deletions, 200 *D. melanogaster* genomes with novel random single nucleotide and deletion polymorphisms were generated computationally. Two different classes of genomes were created, 100 with deletions 1–10 bp in size, and 100 with deletions 1–1,000 bp in size (Table S3). Synthetic reads were generated based on these genomes and aligned and analyzed using the same steps as the experimental samples. A total of 713 synthetic deletions were generated, with 339 1–10 bp deletions and 374 1–1,000 bp deletions. SAMtools identified 86% of 1–10 bp deletions on the *X, 2*^*nd*^, and *3*^*rd*^ chromosomes. Pindel recovered 57% of synthetic 1–1000 bp deletions (213 of the 374) with the highest fraction of deletions recovered on chromosome *2L* (72%) and the fewest on chromosome *2R* (44%) (Table S7). These models indicate that had deletions occurred at a rate higher than observed in wild type, the additional small and large deletion polymorphisms created by error-prone repair mechanisms should have been detected in *c(3)G* or *corolla* females. Taken together, it is most likely that DSBs in SC-deficient flies are repaired by a higher-fidelity repair process, such as canonical NHEJ or sister chromatid repair. When considering the decreased recovery of ring chromosomes in *c(3)G* mutants (Sandler 1965), the simplest explanation for DSB repair in SC-deficient females is by sister chromatid repair.

### D*e novo* transposable element insertions are more frequent in *c(3)G*^*68*^ homozygous females

While the SC is essential for repair of DSBs as CO and NCOGCs, it is unknown if the SC regulates other molecular events such as the movement of TEs. Absence of the yeast *c(3)G* homolog *Zip1* has been shown to result in a decreased insertion rate of the retrotransposon Ty1, suggesting there is a role for the SC in TE movement (Dakshinamurthy *et al.* 2010). The rate at which TE movement occurs during Drosophila meiosis is unknown, therefore to determine the baseline transposition rate, we utilized previously published data and identified forty-four novel TE insertions from the *X, 2*^*nd*^ and *3*^*rd*^ chromosomes from 196 wild-type individuals (Miller *et al.* 2016b) (Figure 2, Table S8). In this dataset a single novel insertion on the *4*^*th*^ chromosome was observed but is not included in the rate calculations (Table S8). Seven of the 44 insertions occurred close enough to a polymorphism to confirm through linkage that they could only have been maternally inherited. For example, male cs13.13 carries a novel *roo* insertion on the *w*^*1118*^ *X* chromosome that is not seen in the 11 other male siblings that also inherited the *w*^*1118*^ haplotype from the same female. It is not possible to definitively determine which parent the remaining 37 events were inherited from due to low SNP density. Using these data, a per arm rate of *de novo* euchromatic TE insertion can be estimated as 0.18 insertions per arm per meiosis [(44 events x 4 haploid meiotic products) / (196 meiosis * 5 arms)], meaning that while a novel TE insertion occurs in approximately 1 in every 5 meioses, it would only be recovered in approximately every 1 in 20 progeny.

**Figure 2.**
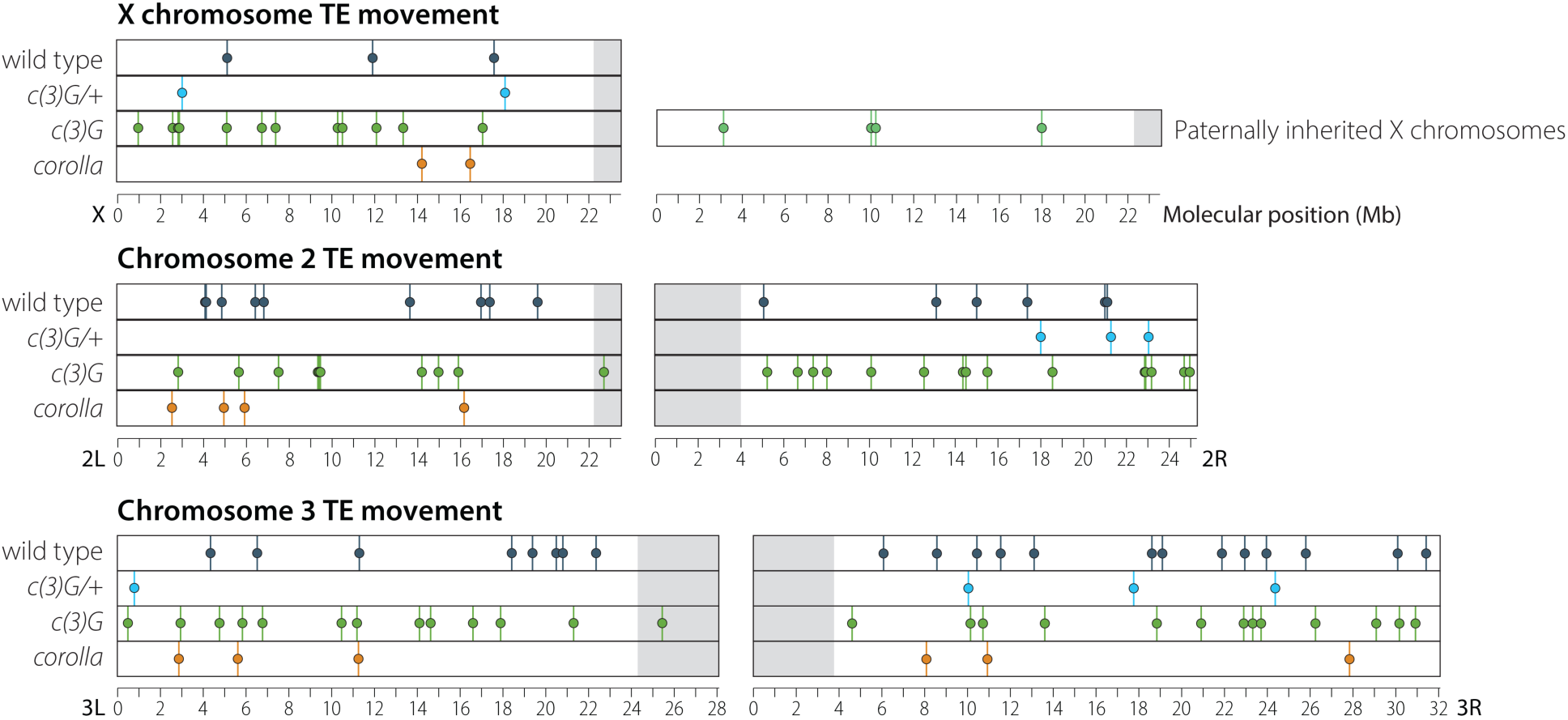
Novel TE insertion positions identified after a single round of meiosis for all four classes of offspring analyzed in this study. Details about insertion position and class of TE inserted can be found in Table S8.

This same approach was then applied to the *X, 2*^*nd*^, and *3*^*rd*^ chromosomes from *c(3)G/+* and *c(3)G* offspring. In the 40 *c(3)G/+* offspring, 9 novel insertion events were identified—2 on the *X* chromosome, 3 on chromosome *2R*, 1 on chromosome *3L*, and 3 on chromosome *3R* (Figure 2, Table S8). Recovery of 9 novel insertion events in 40 individuals when surveying 5 chromosome arms gives a per arm *de novo* rate of transposition of 0.18 insertions per arm per meiosis, identical to the rate observed above in wild type.

In *c(3)G* homozygotes, *de novo* transposition events were identified on all *2*^*nd*^ and *3*^*rd*^ chromosomes as well as the 68 maternally inherited *X* chromosomes from the 93 non-intersex males; the 25 paternally inherited *X* chromosomes were analyzed separately. For maternally inherited chromosomes, 64 novel transposition events were identified—12 on the *X* chromosome, 10 on *2L*, 15 on *2R*, 13 on *3L*, and 14 on *3R* (Figure 2, Table S8). Analysis of sibling *X* chromosome haplotypes confirmed that all 12 maternally inherited *X* chromosome insertions were *de novo*. Among the 25 paternally inherited X chromosomes, 4 *de novo* TE insertions were observed. Considering only maternally inherited chromosomes, the per arm rate of novel transposon insertion events in *c(3)G* male offspring was 0.70 for the *X, 2*^*nd*^, and *3*^*rd*^ chromosomes, significantly higher than wild type (p < 0.001, Chi-square test). The rate of *de novo* transposition events for paternally inherited *X* chromosomes was 0.64, similar to the rate of 0.70 observed in maternally derived chromosomes and also significantly higher than wild type (p = 0.002, Chi-square test).

To help delineate whether the increase in *de novo* transposition events is a general property of SC-deficient females or specific to *c(3)G*^*68*^ homozygous females, novel TE insertions were identified in offspring from the previously described *corolla* mutant females. Using the same approach as above, 12 *de novo* transposon insertions were identified on the *X, 2*^*nd*^, and *3*^*rd*^ chromosomes of 50 individual offspring (Figure 2, Table S8). Of the 12 novel insertions identified, none occurred on the *X* in a male with a paternally inherited *X* chromosome, and one occurred on the *X* chromosome of an *XXY* female carrying two maternally inherited *X* chromosomes. The distribution of events was similar to that observed in wild type. These 12 events were recovered from all five chromosome arms, giving a rate of 0.19 insertions per arm per meiosis in *corolla* mutants, similar to the rate of 0.18 observed in both wild type and *c(3)G*^*68*^ heterozygous females but significantly less than the observed *de novo* TE insertion rate in *c(3)G*^*68*^ homozygous females. This suggests that the increase in *de novo* TE insertions may not be a general property of SC-deficient mutants, but is specific to *c(3)G*^*68*^ homozygotes.

The increased rate of *de novo* transposition in an SC mutant may provide new clues to the role SC components might play in facilitating or preventing the movement of TEs. The observed rate of novel TE insertions in this study was significantly higher in offspring from *c(3)G*^*68*^ females when compared to the other three classes of progeny studied: wild type, *c(3)G*^*68*^ heterozygotes, and *corolla*^*129*^ homozygotes. Somewhat surprisingly, the elevated rate in *c(3)G*^*68*^ maternally derived chromosomes was similar to the rate from paternally derived *X* chromosomes, which came from males with two wild-type copies of *c(3)G*. That the rate of insertion was equal in both maternally and paternally inherited chromosomes was unexpected and suggests that SC proteins may play previously unappreciated roles during both male and female meiosis.

The increased rate of novel TE insertions in *c(3)G*^*68*^ mutants could be explained by a model in which C(3)G prevents mobilized TEs from inserting into genomic DNA. In the absence of C(3)G a greater number of TEs may be available to insert into nuclear DNA. A higher number of active TEs may also explain why the rate of TE insertions was similar on *X* chromosomes derived from wild type males, but would require that TE insertions occur post-fertilization.

Another possible explanation for the increased rate of TE insertion in *c(3)G* females is differences in genetic background leading to an increased rate of transposition (Kidwell *et al.* 1977). Previous studies have reported “bursts” of TE insertions from a specific TE class and attributed the observation to differences in genetic background (Pasyukova and Nuzhdin 1993; Page *et al.* 2007; Guerreiro 2011). It is worth noting that 20 novel insertions in the homozygous *c(3)G*^*68*^ dataset were *doc* elements, which does raise the possibility of differences in genetic background leading to an increased rate of transposition. Although, the genetic background in these experiments was somewhat controlled as females both heterozygous and homozygous for *c(3)G*^*68*^ were heterozygous for the same *w*^*1118*^ and *Canton-S X* and *2*^*nd*^ chromosomes, which were from the same stocks used in the wild-type experiment (Miller *et al.* 2016b). These two stocks differed in that females heterozygous for *c(3)G*^*68*^ carried one copy of a *w*^*1118*^ *3*^*rd*^ chromosome, while those homozygous for *c(3)G*^*68*^ did not. The background of *corolla* mutants was not controlled. Thus, while this may reduce the likelihood of genetic background contributing to the elevated TE insertion rate, it does not completely eliminate it.

A unifying explanation may be that *c(3)G* itself plays a previously unappreciated role in the prevention of TE movement and that this is separate from the role, if any, played by fully functional SC. Despite not building SC, *D. melanogaster* males express *c(3)G*, and other SC genes during meiosis (Brown *et al.* 2014). The reason for this is unclear, but it could be that C(3)G modulates TE movement during both male and female meiosis. This type of role could help explain why SC components are among the most rapidly evolving of all genes (Fraune *et al.* 2012; Hemmer and Blumenstiel 2016) and could be clarified with experimental approaches that delineate the rate of TE insertions in male and female meiosis, control for genetic background, and use sequencing technologies which more reliably identify TE insertions in the genome.

### *De novo* copy-number variation occurs in the absence of full-length SC

Copy number variation is a significant source of genetic variability within populations (Kaminker *et al.* 2002; Lee and Langley 2010). CNVs may be beneficial or deleterious to an individual and may involve a large or small number of genes. Previous studies in *D. melanogaster* have revealed surprisingly high rates of *de novo* CNV both in single offspring or shared among several siblings (Watanabe *et al.* 2009; Miller *et al.* 2016b). In wild type CNVs frequently formed between sister chromatids and were flanked by transposable elements, suggesting that TEs may play a key role in *de novo* CNV formation (Miller *et al.* 2016b). Whether between sister or between homolog CNV formation is dependent on functional SC is unclear.

Large CNVs were identified by plotting depth of coverage for individual chromosome arms. Plots for all chromosome arms for all individuals from *c(3)G*^*68*^/+, *c(3)G*^*68*^, or *corolla*^*129*^ females were generated and revealed 5 total events. No *de novo* CNV events were observed in offspring from *c(3)G*^*68*^/+ females, 4 events were recovered in offspring from *c(3)G*^*68*^ homozygous females, and 1 event in offspring of *corolla*^*129*^ homozygotes (Figure 3, Table S9). Among the 4 events recovered from *c(3)G*^*68*^ homozygous females one was shared among multiple siblings—a 223 kb deletion of chromosome *2R* involving 27 genes, which was recovered from 14 males from females 4, 5, and 6 (Figure 3A). That this event was observed in individuals from multiple crosses of individual male and female makes it likely that the deletion occurred at least two generations prior and did not significantly reduce the fitness of those individuals carrying it.

**Figure 3:**
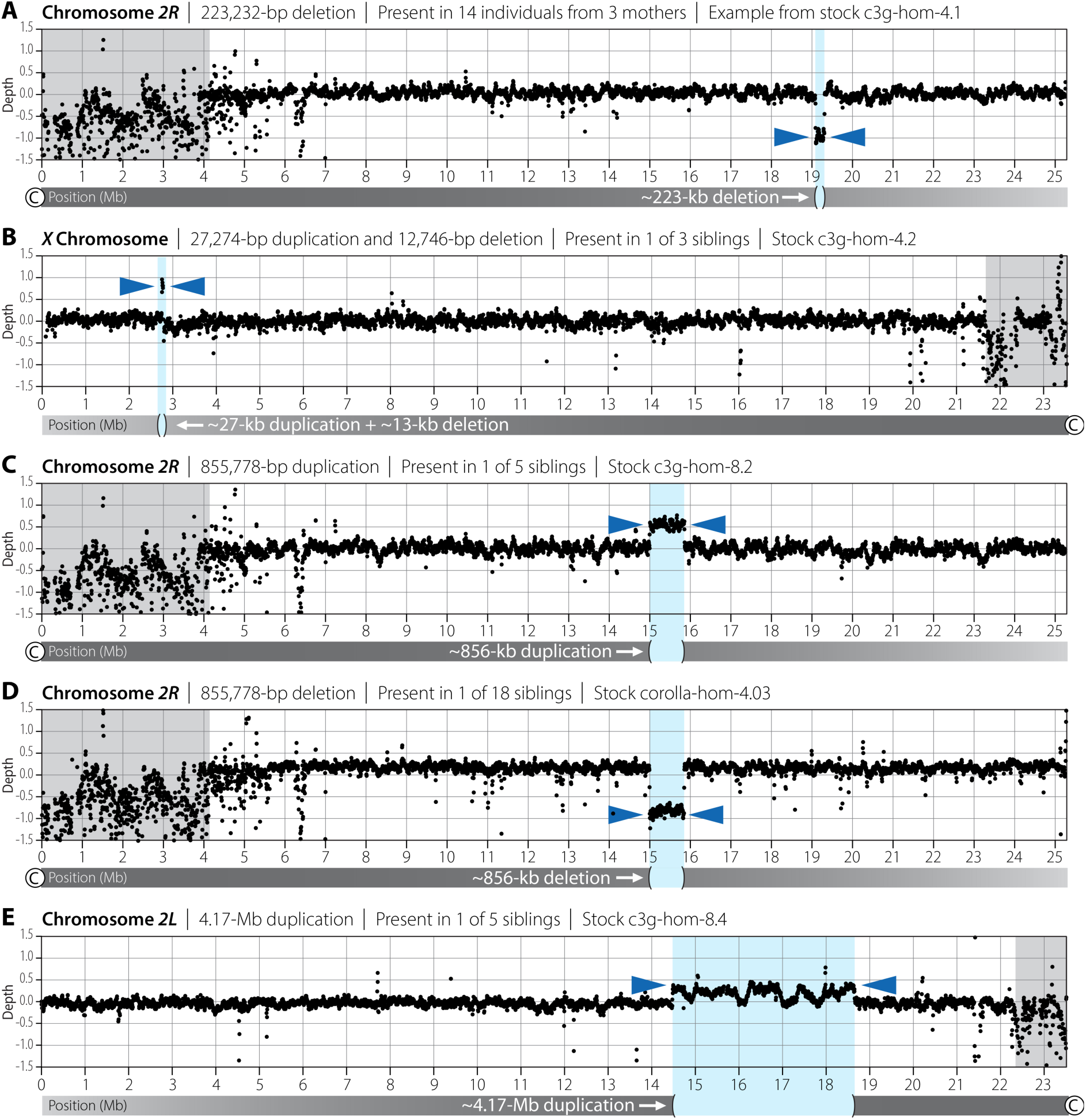
Copy-number variants recovered in this study; details in Table S9. **A.** 223-kb deletion shared among 14 males from 3 different females that likely occurred at least two generations prior. **B.** A complex 27-kb duplication and 13-kb deletion at the *w* locus that was recovered in a single offspring and is likely to be de novo based on sibling haplotypes lacking the rearrangement. **C–D.** An 856-kb duplication identified in a single male from a *c(3)G*^*68*^ homozygous female has identical start and end coordinates as an 856-kb deletion recovered in a single male from a *corolla*^*129*^homozygous female and is identical to an 856-kb duplication recovered in a single male from a wild-type female from a prior study (Miller *et al.* 2016b). **E.** A large 4.17-Mb duplication observed in a single individual that is likely mosaic based on its lower log(2) ratio of 0.25.

The remaining CNVs recovered were only observed in single individuals and, based on haplotype analysis, were likely *de novo* events. One event, a complex *de novo* CNV involving both a deletion and a duplication was identified at the *w* locus on chromosome *X* in a single male from a *c(3)G*^*68*^ homozygous female (Figure 3B). This was an event mediated by unequal crossing over between *Roo* elements. Previous work describing ectopic recombination in *D. melanogaster* focused on unequal exchange between *Roo* elements at the *w* locus, similar to the event observed here (Goldberg *et al.* 1983).

Two *de novo* CNVs, one duplication and one deletion, occurred at the exact same position on chromosome *2R* in two individuals. One of these events, a duplication, was recovered from the offspring of a *c(3)G*^*68*^ homozygous female while a deletion was recovered from the offspring of a *corolla*^*129*^ homozygous female (Figure 3C,D). This 856-kb event includes 107 genes and is flanked on both sides by a hobo element. Remarkably, a CNV with the exact same breakpoints was identified in a previous study of individuals from wild-type females (Miller *et al.* 2016b). All three of these CNVs were *de novo* events validated using the haplotypes of the siblings that did not carry the CNV. All three crosses were between individual males and females and multiple genetic backgrounds are involved (*w*^*1118*^, *Canton-S*, and the undefined *corolla*^*129*^ background) thus these CNVs are not variants segregating at low frequency in the population and are recurrent *de novo* events.

The final CNV observed was a 4.2-Mb duplication on chromosome *2L* not flanked by a TE or low-complexity sequence recovered in a single male from a *c(3)G*^*68*^ homozygous female (Figure 3E). Analysis of read pairs demonstrate that this is a tandem duplication as reads mapping to the proximal end of the duplication are linked to reads mapping to the distal end of the duplication. The log_2_ depth-of-coverage ratio for this interval is 1.25, 0.25 higher than expected for a diploid and less than the log_2_ depth-of-coverage ratio of 1.5 that would be seen in an autosomal duplication occurring before the first mitotic division. Thus, this duplication is present in half the cells in the individual sequenced and likely occurred during the first mitotic division, possibly as a consequence of a re-replication event that was then repaired by recombination between the duplicated segments (Green *et al.* 2010). It is notable that the fly was able to tolerate such a large duplication, involving 513 genes, present in half of all cells. Although the possibility that there was selection against cells carrying the large duplication cannot be excluded, a log_2_ depth-of-coverage ratio of 1.25 does strongly suggest there was limited selection against those cells with the duplication. If selection was acting strongly on these cells the log_2_ ratio would fall below 1.25 and perhaps become undetectable.

The recovery of TE-mediated CNVs in females unable to construct SC demonstrates that these CNVs can occur independently of normal meiotic synapsis and DSB formation, perhaps depending only on the presence of a chromosome axis. It is also possible these events may occur during mitosis. That two different TE-mediated CNV events, a deletion and a duplication, were recovered at the exact same coordinates in two different genetic backgrounds as a duplication observed in wild-type individuals was surprising and suggests that the rate of CNV formation is not uniform across the genome. As would be expected in mutants with defective homologous chromosome pairing, all four TE-mediated CNV events recovered appear, based on allele frequency and TE positioning, to be events between sister chromatids and not between homologous chromosomes.

## ACKNOWLEDGEMENTS

I would like to thank Angela Miller for editorial and figure preparation assistance; the Stowers Institute Molecular Biology core for expert assistance with DNA sequencing; Nazanin Yeganeh Kazemi and Clarissa Smith for assistance with DNA preparation and NCOGC validation; Justin Blumenstiel, Cori Cahoon, and Talia Hatkevich for helpful discussion and comments; and R. Scott Hawley for critical feedback and invaluable mentorship. The DNA sequence data described here were supported by funding to Scott Hawley from the Stowers Institute for Medical Research during my work in his lab.

## SUPPLEMENTAL FIGURES

**Figure S1:** Cross schemes. **A.** Isogenic Canton-S females carrying the loss-of-function mutant *c(3)G*^*68*^ were crossed to isogenic *w*^*1118*^ males. Individual females heterozygous for the *c(3)G*^*68*^ mutant allele were collected and crossed to individual isogenic *w*^*1118*^ males and individual male offspring were collected and sequenced. **B.** Isogenic *Canton-S* females carrying the loss-of-function mutant *c(3)G*^*68*^ were crossed to isogenic *w*^*1118*^ males carrying the loss-of-function mutant *c(3)G*^*68*^. Individual females hemizygous for both mutant alleles were collected and crossed to individual isogenic *w*^*1118*^ males. Individual phenotypically male offspring were then collected and sequenced **C.** Individual females homozygous for *corolla*^*129*^ were crossed to individual male siblings and individual male and female offspring were then collected and sequenced.

**Figure S2:** Log_2_ depth-of-coverage analysis for chromosome *2L*, the *X* chromosome, and the *4*^*th*^ chromosome. Wild-type data from Miller *et al*. **(Miller *et al.* 2016b)**, genotypes are given for each individual. This analysis uncovered three intersex males and six individuals with an extra copy of chromosome *4*. One of the intersex males and one of the XY males are also mosaic for loss of a *4*^*th*^ chromosome, meaning some of their cells have 3 *4*^*th*^ chromosomes while others have 2 *4*^*th*^ chromosomes. The log_2_ differences for the *X* and *4*^*th*^ chromosomes use chromosome *2L* as the basis of their log_2_ ratio calculation (Table S1).

**Figure S3:** Three intersex males were based on depth of coverage and their autosomal allele frequency. A heterozygous male should have a 50%/50% *w*^*1118*^/Canton-S allele frequency for the *2*^*nd*^ chromosome, and because they are hemizygous for the *X* chromosome, a 100% allele frequency for either *Canton-S* or *w*^*1118*^ SNPs along the *X* chromosome. These three males carry a 50% *w*^*1118*^/Canton-S allele frequency for the *X*, suggesting that they carry two distinct *X* chromosomes. They also carry a 67%/33% *w*^*1118*^/Canton-S allele frequency for both arms of the *2*^*nd*^ chromosome, with 67% of the SNPs from the *w*^*1118*^ stock, and 33% of the SNPs from the Canton-S genome—evidence for the presence of three *2*^*nd*^ chromosomes.

## SUPPLEMENTAL TABLES

**Table S1:** Details for each individual sequenced in this project including maternal genotype, barcode, and depth of coverage for each chromosome arm.

**Table S2:** Primers used to check gene conversions in males from c3g homozygous mothers.

**Table S3:** List of random deletions generated per genome in order to test the deletion identification pipeline.

**Table S4:** COs recovered in this study.

**Table S5:** NCOGCs recovered in this study.

**Table S6:** Novel deletions identified in all four classes of progeny used in this study.

**Table S7:** Summary of recovery using Pindel of deletions 1-1000bp in size for computationally generated genomes.

**Table S8:** Details of de novo transposable element insertion events.

**Table S9:** Details of one complex and four simple copy-number variants recovered in this study.

